# Deletion of RsmE 16S rRNA methyltransferase leads to low level increase in aminoglycoside resistance in *Mycobacterium smegmatis*

**DOI:** 10.1101/2020.01.15.907279

**Authors:** Shipra Bijpuria, Rakesh Sharma, Bhupesh Taneja

## Abstract

Owing to its central role in cellular function, ribosome is one of the most common targets of antibiotics in bacteria. Mutations in *rrs* gene, ribosomal protein genes, methyltransfersases or drug influx/efflux are often found to overcome the drug response. Despite modulation of methylation pattern in the ribosome through mutations in the methyltransferases as one of key modulators of drug response, *rsmG (gidB)* is the only conserved methyltransferase associated with low-level drug resistance in large number of mycobacterial isolates. Here, we present the first evidence of association of methylation by mycobacterial RsmE, that methylates U1498 of 16S rRNA, with low levels of drug resistance. Deletion of the RsmE-homolog of *Mycobacterium smegmatis* leads to at least two-fold increase in the inhibitory concentration of aminoglycosides that bind in the decoding center proximal to U1498 in the 30S subunit. The change in inhibitory concentrations was highly specific and does not show any cross-resistance to drugs of other classes. Surprisingly, Rv2372c, the RsmE-homolog of *Mycobacterium tuberculosis* has the largest number of mutations among conserved ribosomal methyltransfersases, after *gidB*, highlighting the role of mutations in the RsmE methyltransferase as a key emerging mechanism of drug resistance in clinical strains of *M. tuberculosis*. Our work underlies the association of methylation by the RsmE-homolog with drug resistance and lays the groundwork to tackle this emerging mechanism of drug resistane in mycobacteria.

## Introduction

*M. tuberculosis* is one of the most prevalent pathogens and responsible for largest number of deaths by a single infectious pathogen, annually, worldwide (WHO 2018). Treatment involves multidrug therapy with the four frontline drugs for extended periods of time (Sotgiu *et al.*, 2015). However, a rapid increase in resistance towards the first line of drugs, over the last few years, has led to an increased dependence on a prolonged and complex treatment that includes the second line of drugs, namely, fluoroquinolones and injectable aminoglycosides (kanamycin, amikacin, capreomycin) (Sotgiu *et al.*, 2015; WHO 2018). Any resistance towards the second line of drugs compounds the problem towards extensively drug resistant disease (XDR-TB). Resistance to fluoroquinolones may occur due to mutations in gyrases (Nosova *et al.*, 2013; Zhang and Yew 2015), to kanamycin by overexpression of *eis* drug-modifying enzyme (Sowajassatakul *et al.*, 2014; Kambli *et al.*, 2016) or of efflux pumps (Louw *et al.*, 2009; Reeves *et al.*, 2013), and to other aminoglycosides by modification of target binding site through mutations in *rrs* (Gygli *et al.*, 2017; Hameed *et al.*, 2018), ribosomal protein genes (Finken *et al.*, 1993) or changes in rRNA methylation pattern (Buriánková *et al.*, 2004; Maus *et al.*, 2005; Okamoto *et al.*, 2007).

Methylations in ribosomal RNA are brought about by highly specific rRNA methyltransferases. The complete set of conserved ribosomal methyltransferases has been identified in *E. coli* and indicates ten conserved methyltransferase members for the 30S subunit (Golovina *et al.*, 2012). These rRNA methyltransferases are known to modulate the structure of ribosome by addition of methyl group to key nucleotides in the decoding center of ribosome. Mutation in some of the rRNA methyltransferases is associated with altered function of the ribosome as well as with drug resistance. For instance, mutations in *gidB*, the RsmG 16S rRNA methyltransferase has been reported to be associated with low level resistance towards streptomycin and leads to at least twofold increase in MIC to streptomycin in *E. coli* and *M. tuberculosis* (Okamoto *et al.*, 2007; Wong *et al.*, 2011). Mutations in *tlyA*, that methylates C1920 in 23S rRNA and C1409 in 16S rRNA, are linked to resistance to capreomycin in *M. tuberculosis* (Maus *et al.*, 2005). Despite key roles of methylated nucleotides in the decoding center, Rv3919c (RsmG/GidB, Wong *et al.*, 2011), Rv2966c (RsmD, Kumar *et al.*, 2011) and Rv2372c (RsmE, Kumar *et al.*, 2014) are the only conserved 30S rRNA methyltransferases identified in *M. tuberculosis*. Moreover, the role of these methyltransferases in drug resistance, apart from *gidB*, remains poorly characterized.

We have earlier identified Rv2372c as a RsmE homolog of *M. tuberculosis* that methylates U1498 in helix 44 of 30S subunit of ribosome in a highly specific manner (Kumar *et al.*, 2014). U1498 is one of the conserved methylated nucleotides present in the decoding center of the ribosome that together form a compact hydrophobic cage around the anticodon stem loop structures at the P-site and monitor the codon-anticodon interactions (Korostelev *et al.*, 2006; Selmer 2006). However, deletion of *rsmE* had no significant effect on growth and survival of *E. coli* (Basturea 2006). Apart from its key roles in ribosomal function and fidelity, U1498 also likely to affect the response to antibiotics, as U1498 lies in the vicinity of binding site of various aminoglycosides as seen in the crystal structures of ribosome-antibiotic complexes (Borovinskaya *et al.*, 2007, 2008; Demirci *et al.*, 2013; Cocozaki *et al.*, 2016). A mutation of U1498C is among three mutations that confer different levels of hygromycin resistance in *M. smegmatis in vitro* (Pfister *et al.*, 2003). However, this mutation has not been reported in clinical mycobacterial strains. Moreover, the association of methylation at U1498 has not been previously investigated with altered drug response in mycobacteria.

Given the association of antibiotic response with methylated nucleotides in the decoding center, and the key role of helix 44/ U1498 in the P-site of ribosome, here, we investigate the role of methylation by mycobacterial RsmE with antibiotic response by generating a deletion mutant of the RsmE homolog in *M. smegmatis*. Growth of wild-type, knockout and complemented strain was analyzed with varying concentration of ribosomal- and non-ribosomal-targeting drugs and showed at least two-fold increase in the inhibitory concentration of several antibiotics. Moreover, Rv2372c, the RsmE-homolog of *M. tuberculosis* was found to harbor the largest number of mutations among conserved ribosomal methyltransfersases, apart from *gidB*, suggesting a possible correlation of RsmE mutations with drug response in clinical strains. In this work, we show U1498 methylation is conserved in mycobacteria and present the first evidence of association of U1498 methylation with modulation of antibiotic response in any bacteria.

## Methods

### Mutational analysis of Rv2372c in pathogenic *M. tuberculosis* strains

Whole genome sequencing data for available 9745 strains of *M. tuberculosis* were extracted from PATRIC database (www.patricbrc.org, PATRIC March 2019 Release). Rv1010, Rv1407, Rv2966c, Rv2372c, Rv3919c, Rv2165 and Rv1003 were identified as homologs of conserved 16S rRNA methyltransferases of *E. coli* on the basis of sequence similarity and selected for further analysis. Non-synonymous mutations in the mycobacterial 16S rRNA methyltransferases were identified from genome data of all the strains, with *M. tuberculosis* H37Rv as the reference strain.

### Bacterial strains and growth conditions

*M. smegmatis* mc^2^ 155 (Ms-WT) or *RsmE*-depleted strains (Ms-ΔRsmE) were grown in Difco Middlebrook 7H9 broth or on 7H10 agar plates, supplemented with 0.05% Tween 80, 0.25% glycerol and 0.4% glucose. Growth profiles were obtained by monitoring growth at 600 nm at 37 °C with continuous shaking using the BioScreen growth curve analyzer and plots were generated. The strains, plasmids and the primers used in the study are listed in Table 1.

**Table 1:**
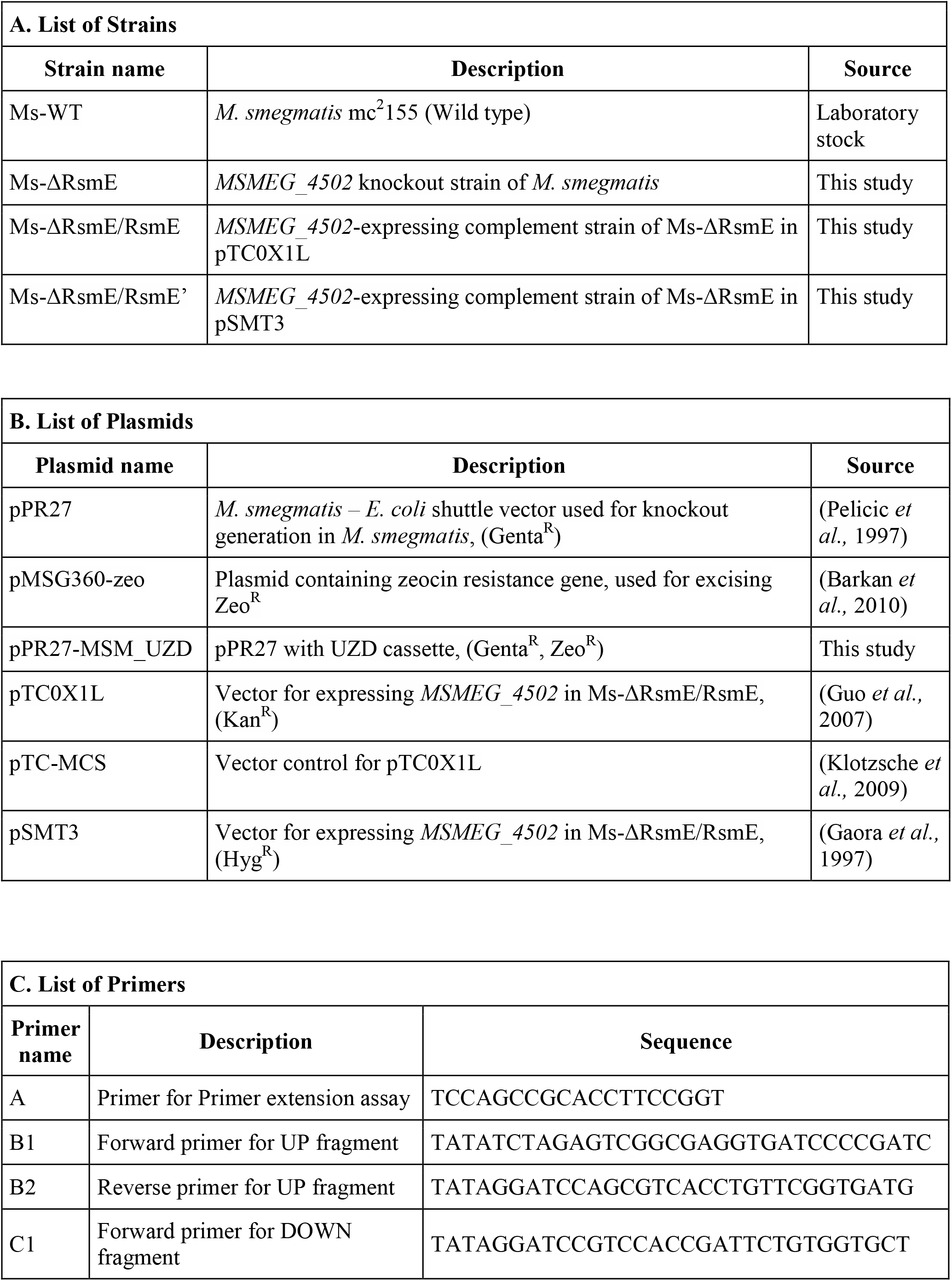
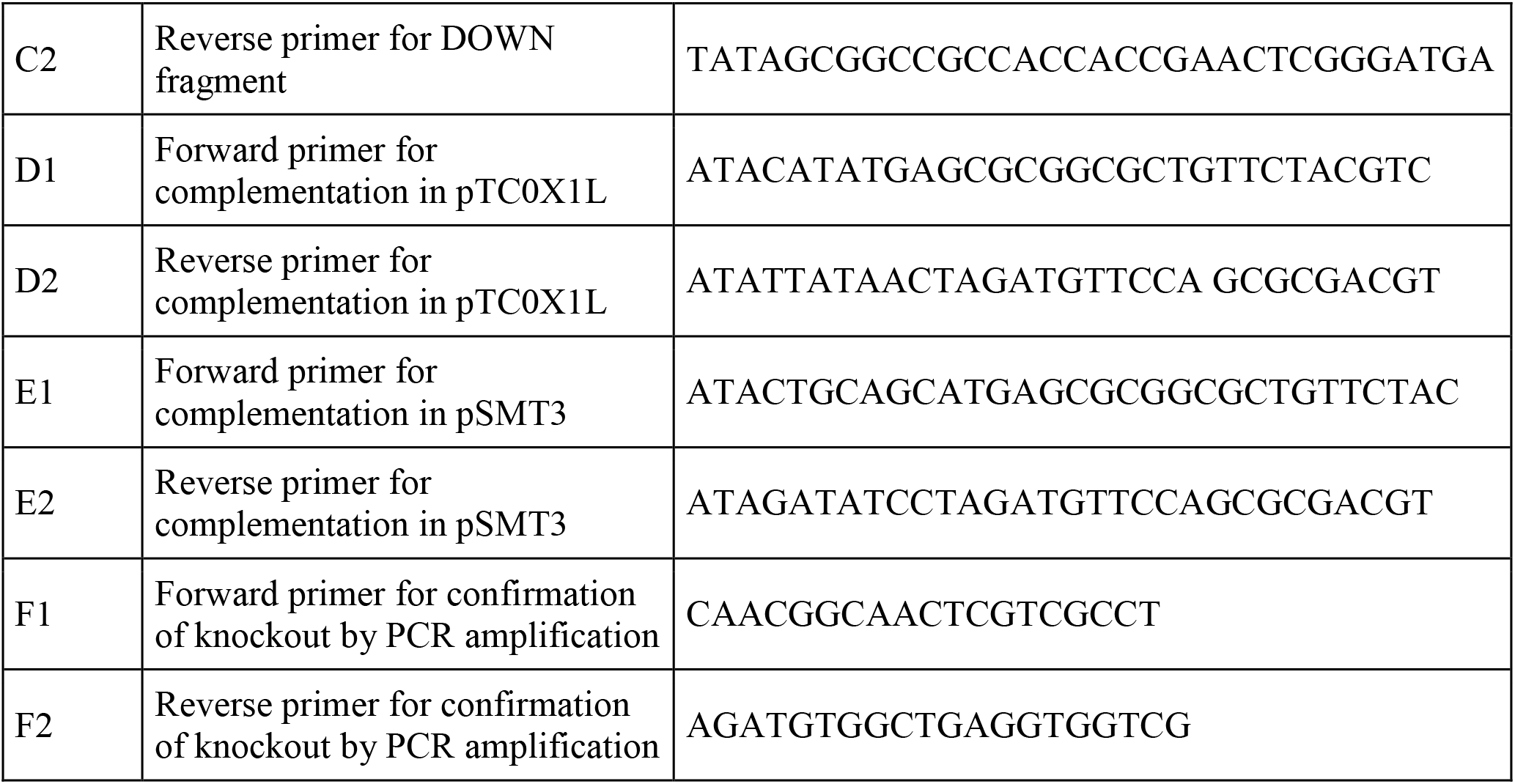
Summary of strains, plasmids and primers used in the study

### Construction of *M. smegmatis* knockout and complementation

Rv2372c was earlier identified as the RsmE ortholog in *M. tuberculosis* (Kumar *et al.*, 2014). A knockout of *MSMEG_4502*, the sequence homolog of *Rv2372c* in *M. smegmatis*, was generated by a suicidal vector strategy using pPR27, as described earlier (Pelicic *et al.*, 1996; Pelicic *et al.*, 1997). Briefly, for the targeted gene knockout, the 5’ and 3’ ends of *MSMEG_4502* and its flanking sequences were amplified to generate ‘UP’ (813 bp) and ‘DOWN’ (823 bp) fragments. A zeocin resistance gene (657 bp) excised from pMSG360-zeo (Barkan *et al.*, 2010)) was introduced between them to generate UZD (UP-Zeocin-DOWN) deletion cassette. The UZD deletion cassette was then cloned between XbaI and NotI sites in *M. smegmatis – E. coli* shuttle vector pPR27 yielding plasmid pPR27-MSM_UZD. Plasmid pPR27-MSM_UZD was then transformed into *M. smegmatis* mc^2^ 155 via electroporation, to generate the knockout by homologous recombination method.

The pPR27-MSM_UZD transformants of *M. smegmatis* were selected on 7H10 agar plates containing 10 μg/ml gentamicin and 40 μg/ml zeocin. Knockout strains were selected by growing at 42 °C to promote loss of the plasmid as it has a temperature sensitive origin of replication, Ori_Myco_TS. The knockouts were screened by a two-step selection method. Firstly, the mutants were grown at 42 °C on 7H10 plates along with 10% sucrose for positive selection of knockouts that have lost *sacB* gene. Mutants were further screened through negative selection by growing on 7H10 plates containing 10 μg/ml gentamicin. The gene knockout of *MSMEG_4502* was finally confirmed by PCR for incorporation of UZD disruption cassette and the knockout strain designated as Ms-ΔRsmE.

For complementation of the knockout strain, *MSMEG_4502* was cloned in pTC0X1L (Guo *et al.*, 2007; Klotzsche *et al.*, 2009) as well as in pSMT3 expression vector (Gaora *et al.*, 1997) and designated as Ms-ΔRsmE/RsmE or Ms-ΔRsmE/RsmE’, respectively. Ms-ΔRsmE/RsmE’ was used for complementation when the response with kanamycin and neomycin was tested. pTC-MCS (Klotzsche *et al.*, 2009) or pSMT3 were used as parent vector controls along with indicated antibiotics for both wild type or knockout strain.

### *In-vivo* methylation assay

To test the activity of MSMEG_4502 in *M. smegmatis* cells, primer extension assay was carried out (Kumar *et al.*, 2014). Briefly, total RNA was isolated from the *M. smegmatis* strains and 16S rRNA was gel purified from the total RNA. A primer was synthesized, complementary to the sequence 1497 to 1515 of 16S rRNA of *M. smegmatis* (corresponding to 1513 to 1531 of *E. coli* 16S rRNA). The primer was labelled at 5’ end with ^32^P-labelled ATP, using T4 Polynucleotide kinase. Primer extension was done using Verso reverse transcriptase as per the manufacturer’s protocol. The reaction was terminated by heating at 80 °C for 15 minutes and the products were analysed by running on a 15% urea-PAGE. The gel was further imaged by phosphorimaging. The expected size of the product is 33 bp long if U1482 is methylated (1483 bp to 1515 bp). U1482 of *M. smegmatis* 16S rRNA (NCBI Accession No. NC_008596.1:c5029475-5027948) corresponds to U1498 of *E. coli* 16S rRNA (NCBI Accession No. NC_000913.3:4166659-4168200). An oligonucleotide of 33 bp size was *5’* end labelled with ^32^P and used as an indicative size marker.

### Phenotypic Characterisation of knockout strains

For the growth assay, Ms-WT, Ms-ΔRsmE or Ms-ΔRsmE/RsmE were grown till an OD (600 nm) of ~ 1.0 at 37 °C. It was then diluted 1:1000 with fresh media and growth was monitored every 2 hours using the Bioscreen growth analyzer.

Growth assay was also performed to investigate the effect of stress conditions on the growth of knockout. For this, *M. smegmatis* strains were grown in 7H9 media along with varying concentrations of different stress agents, as indicated. Growth under oxidizing conditions was monitored by growing cells in media containing increasing concentrations (0, 0.25, 0.5, 1.25 and 1.5 mM) of hydrogen peroxide while effect of osmotic stress was monitored with increasing concentrations (0, 0.4, 0.6, 0.8 and 1.0 M) of NaCl. Knockout and wild type cells were grown in sodium nitroprusside (0, 10, 20, 30, 40 and 50 mM) to monitor SNP-mediated stress, which is a NO donor and induces nitrosative stress. For acid stress, knockout or wild type strains were grown in 7H9 media at pH 6.0, pH 5.5, pH 5.0 or pH 4.5.

### Drug sensitivity of knockout strain

Drug sensitivity of the knockout strain was tested by microbroth dilution method. Briefly, *M. smegmatis* strains, Ms-WT, Ms-ΔRsmE and Ms-ΔRsmE/RsmE (or Ms-ΔRsmE/RsmE’) were first grown until OD (600nm) ~ 1.0 at 37 °C. The respective strains were then diluted 1:1000 with fresh media and grown with serial dilution of indicated antibiotics added to the culture. Untreated culture and blank media was taken as positive and negative control, respectively. The growth was monitored at 600 nm every 2 hours at 37 °C with continuous shaking in the Bioscreen growth analyzer. The growth curve was plotted using the average value of three replicates. MIC was reported as the lowest drug concentration showing no visible growth in the respective antibiotics.

Antibiotics used in the study belonged to the following three groups: 30S ribosomal-targeting drugs - kanamycin, amikacin, gentamicin, neomycin, paromomycin, tetracycline and streptomycin; 50S ribosomal-targeting drugs - chloramphenicol and erythromycin and non-ribosomal-targeting drugs - moxifloxacin and rifampicin.

## Results

### Generation of *rsmE* knockout of *M. smegmatis*

MSMEG_4502, was identified as the homolog of RsmE of *E. coli* (e-value: 4e^-26)^ and Rv2372c of *M. tuberculosis* (e-value: 3e^-^) on the basis of sequence similarity. Knockout of *MSMEG_4502* in *M. smegmatis* was generated with the help of a suicidal vector strategy (Figure 1A) as described in the Methods section. The mutants were screened and confirmed by PCR (Figure 1B) as well as by DNA sequencing. For confirmation by PCR, primers were designed flanking the UP and DOWN fragments (primer F1 and primer F2, Table 1, respectively). An expected amplified product size of 2535 bp in the knockout (lane 1) as compared to that of 2160 bp in Ms-WT (lane 2), confirms the successful knockout of *MSMEG_4502* in *M. smegmatis* to generate Ms-ΔRsmE.

**Figure 1:**
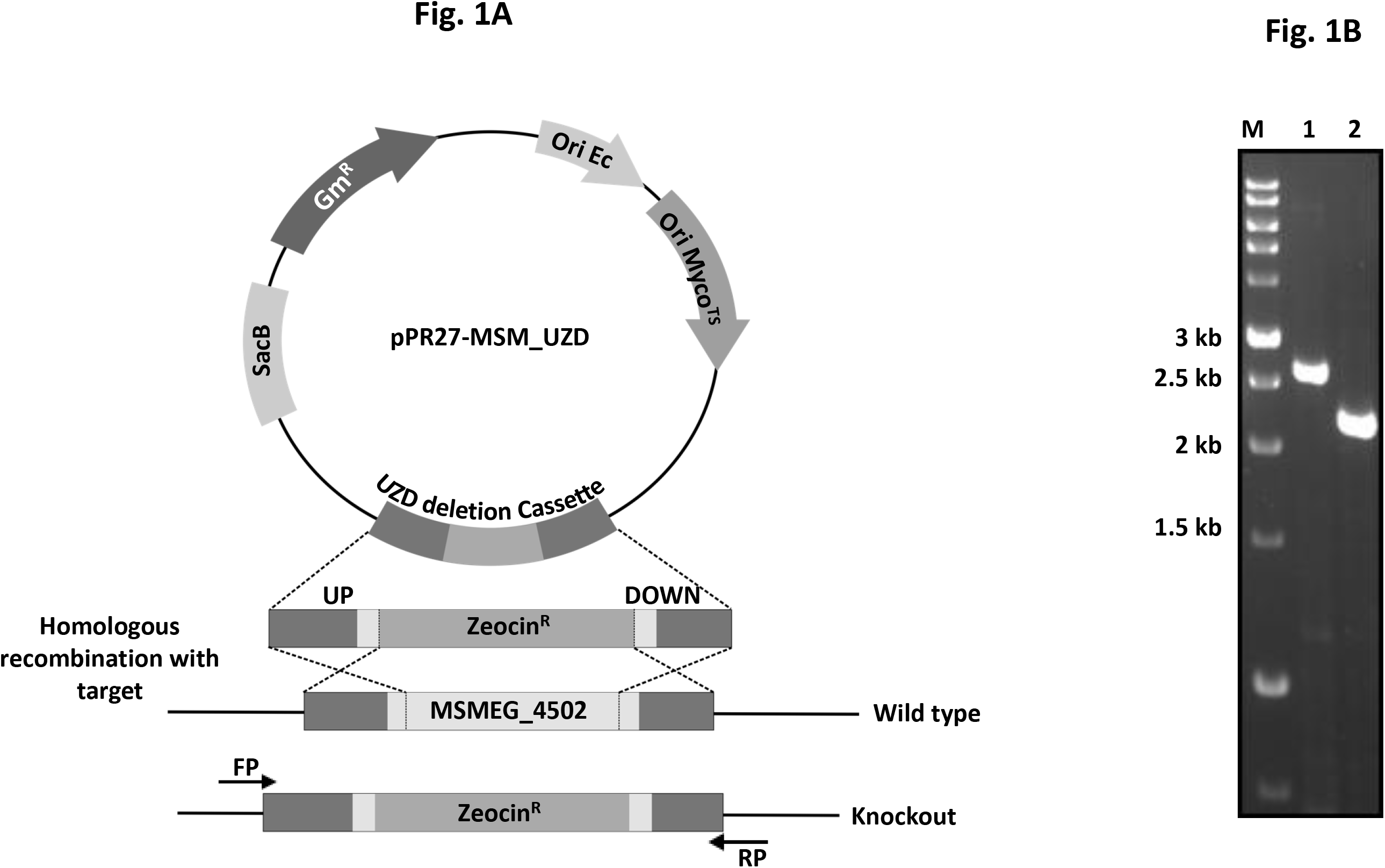
Generation of *M. smegmatis RsmE (MSMEG_4502)* knockout strain. A. A knockout of MSMEG_4502 was generated using suicidal vector strategy with *M. smegmatis - E. coli* shuttle vector pPR27. The upstream fragment (UP) and downstream fragment (DOWN) of the target gene *MSMEG_4502* were cloned in the vector along with Zeocin^R^, yielding pPR27-MSM_UZD. The knockout was generated by homologous recombination and further screening of the transformants. B. The knockouts were confirmed via PCR using the primers flanking the UP and DOWN fragments. An amplified product of 2535 bp is obtained in Ms-△RsmE (Lane 1) and that of 2160 bp in Ms-WT (Lane 2).

### *In-vivo* methylation of 16S rRNA by MSMEG_4502 in *M. smegmatis*

The methylation status at U1498 of 16S rRNA by the mycobacterial homolog was examined by primer extension assay on purified 16S ribosomal RNA extracted from the wild-type (Ms-WT), knockout (Ms-ΔRsmE) and complemented (Ms-ΔRsmE/RsmE) strains. Methylation occurs at N-3 position of U, disrupting hydrogen-bonding with a complemetary A-nucleotide during reverse ß transcription. A primer extension by reverse transcriptase hence stops 3’ to the methylated m^2^ U. As seen in Figure 2, extension of primer (Primer A, Table 1) complementary to 1497 to 1515 by reverse transcriptase stops at U1483 if U1482 is methylated (equivalent to U1498 of *E. coli* 16S rRNA) resulting in a 33 bp product in wild type and complemented strains while reverse transcriptase reads through position 1482 to yield longer products in Ms-ΔRsmE (Figure 2B). Hence, MSMEG_4502 is the RsmE-homolog in *M. smegmatis* that specifically methylates the equivalent m^3^ U1498 position in 16S rRNA in *M. smegmatis.*

**Figure 2:**
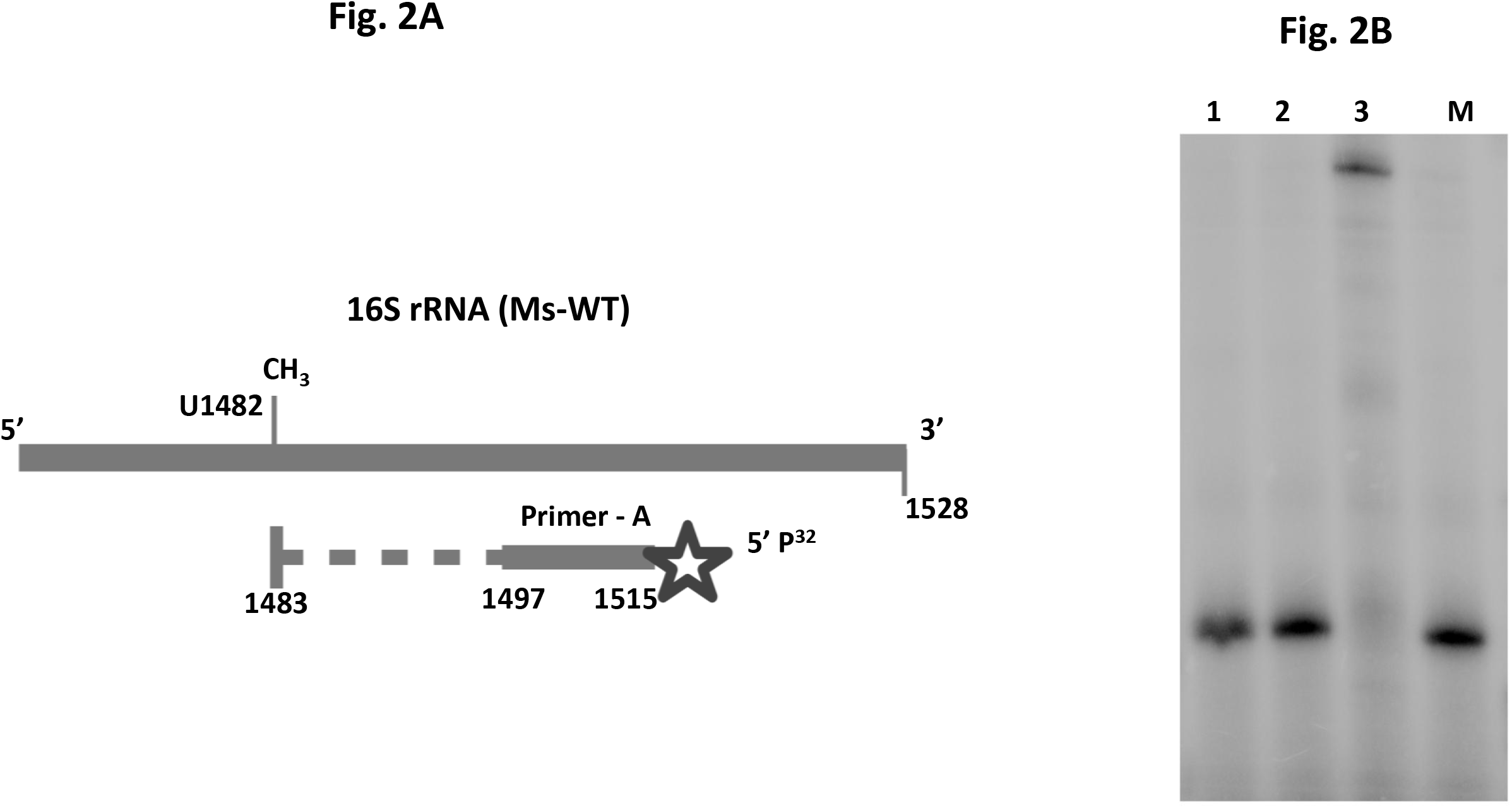
Primer extension assay for m^3^ U1498 methylation in *M. smegmatis*. A. Overall assay design of the primer extension assay. The primer complementary to the region 1497 to 1515 of 16S rRNA of *M. smegmatis* (Primer-A, Table 1) is 5’end-labelled with ^32^P. In the presence of methylation, the reverse transcription is stalled due to disruption of base pairing, giving a truncated product of 33bp, while in the absence of methylation, reverse transcription continues resulting in longer products. All indicated position numberings are with respect to *M. smegmatis* 16S rRNA. *M. smegmatis* U1482 is equivalent to U1498 of *E. coli* 16S rRNA. B. Primer extension assay products from wild type, knockout and complemented strains of *M. smegmatis* were analysed on a 15% urea PAGE. The presence of a band indicates a truncated product of 33bp in Ms-WT (lane 1) and Ms-△RsmE/RsmE (lane 2) due to presence of m^3^ U methylation. (As explained above, *M. smegmatis* U1482 is equivalent to U1498 of *E. coli* 16S rRNA). Absence of methylation in Ms-△RsmE (lane 3) leads to longer products. A^32^P end-labelled 33 base oligonucleotide was used as a marker (M) to indicate the size of the truncated product due to methylation in Ms-WT and Ms-△RsmE/RsmE.

### Effect on growth of Ms-ΔRsmE in the absence of U1498 methylation

*M. smegmatis* strains Ms-WT and Ms-ΔRsmE, with respective empty vector controls, along with Ms-ΔRsmE/RsmE were grown in 7H9 media at 37 °C and growth monitored at OD (600 nm) in the Bioscreen growth analyzer. Ms-ΔRsmE showed slightly slower growth as compared to Ms-WT (Figure 3) that is partially restored in the complemented strain.

**Figure 3:**
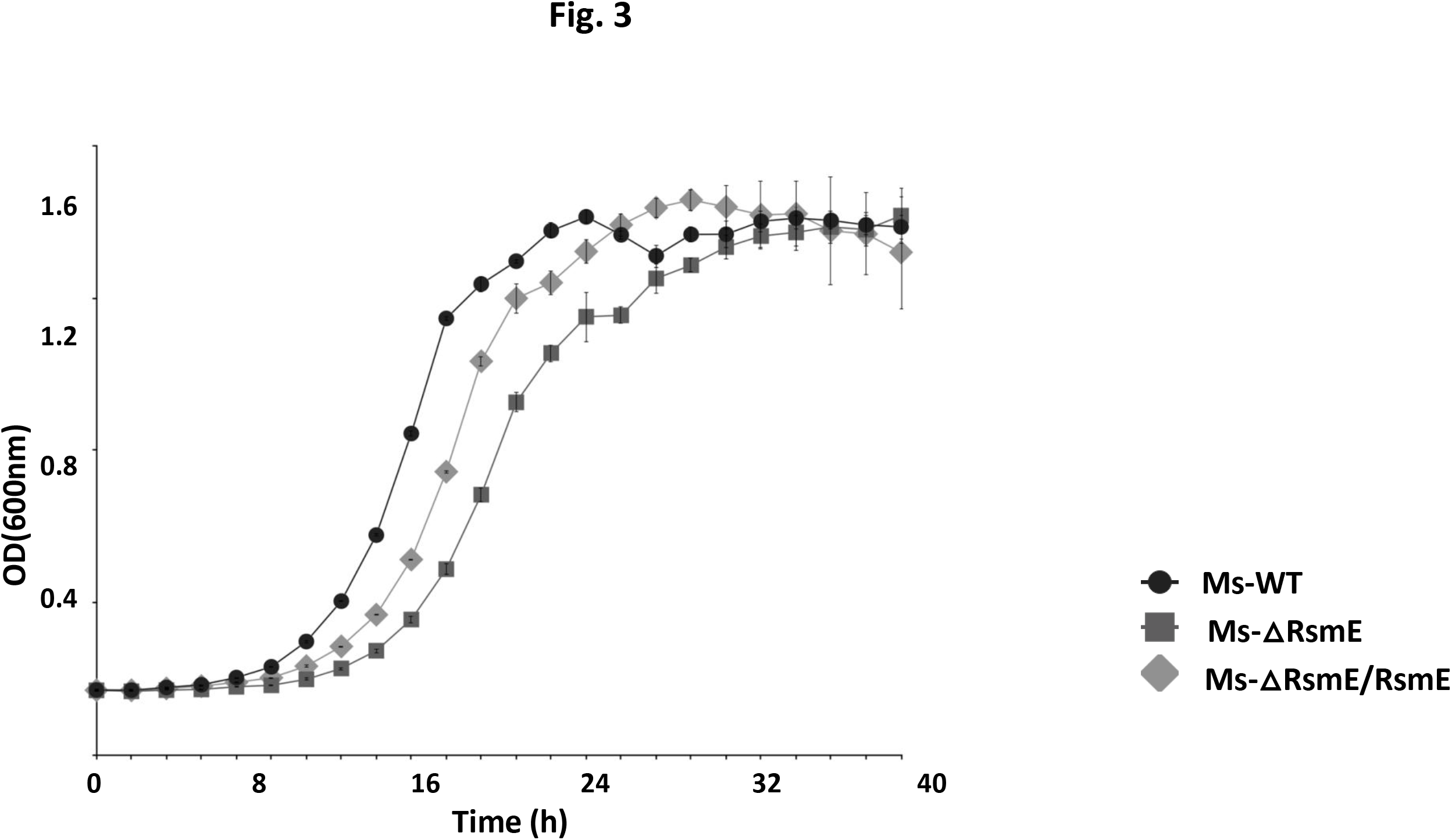
Growth curve of *Mycobacterium smegmatis* strains. The Ms-ΔRsmE knockout strain (squares) shows a slightly extended lag phase compared to Ms-WT strain (circles), which is partially restored, similar to Ms-WT, in the complemented strain Ms-ΔRsmE/RsmE (diamonds).

The growth of Ms-ΔRsmE was also monitored under different stress conditions. Effect of oxidative stress, osmotic stress, SNP induced stress and acid stress on the growth of Ms-WT, Ms-ΔRsmE and Ms-ΔRsmE/RsmE strains were examined. For effect on growth of Ms-ΔRsmE, under oxidative stress, cells were grown in the presence of increasing concentrations of hydrogen peroxide. Deletion of the RsmE-homolog showed no significant difference on growth under oxidative stress conditions and the growth of Ms-ΔRsmE was largely similar to Ms-WT (Figure 4A). To induce osmotic stress to the mycobacterial strains, cells were grown in increasing concentrations of sodium chloride and growth was monitored by OD_600_. Ms-ΔRsmE showed similar growth as Ms-WT under osmotic stress as well (Figure 4B). The growth of Ms-ΔRsmE was slow in response to sodium nitroprusside, a nitrosative stress agent which induces stress by generating NO inside the cells (Figure 4C).

**Figure 4:**
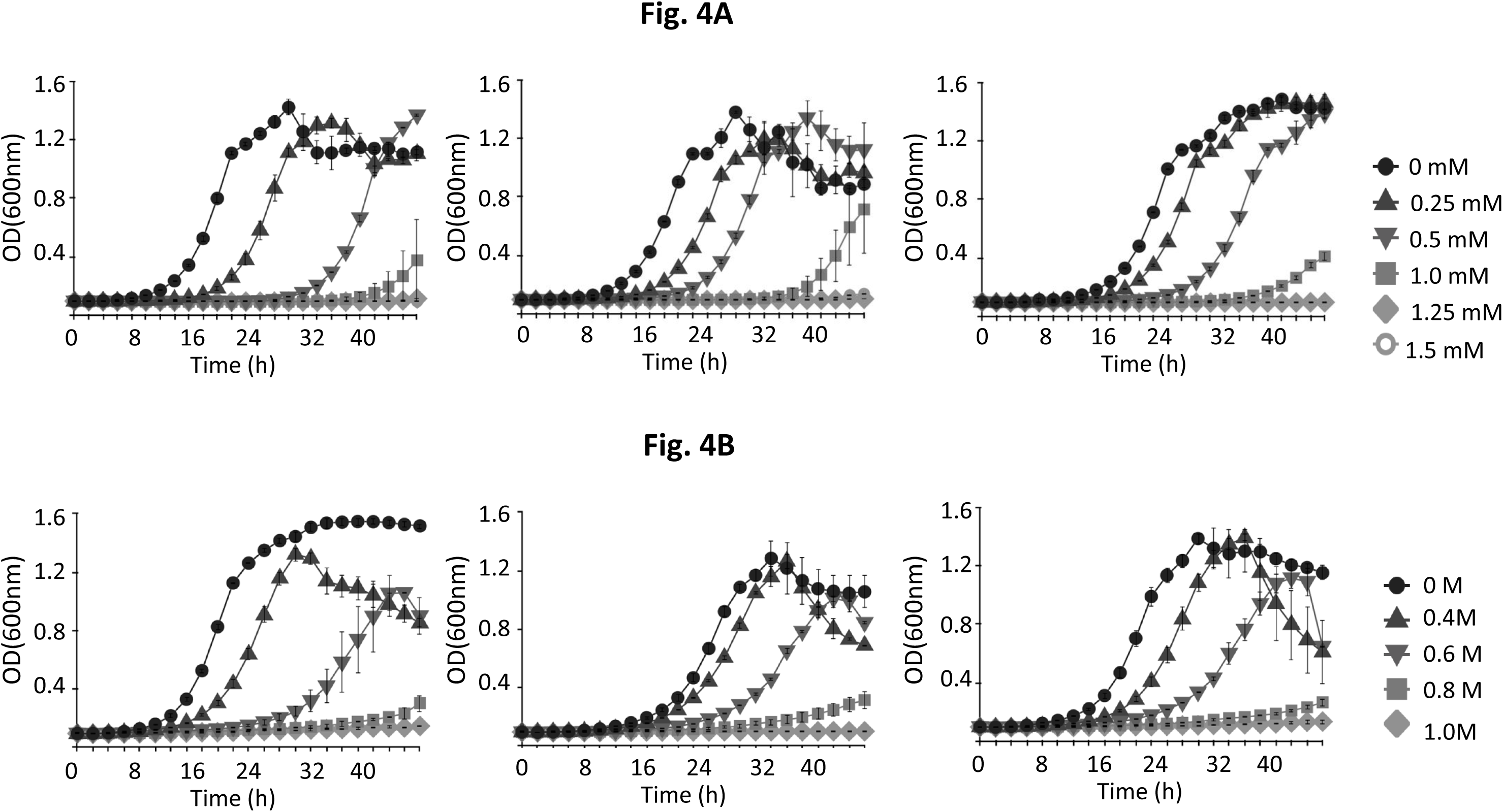
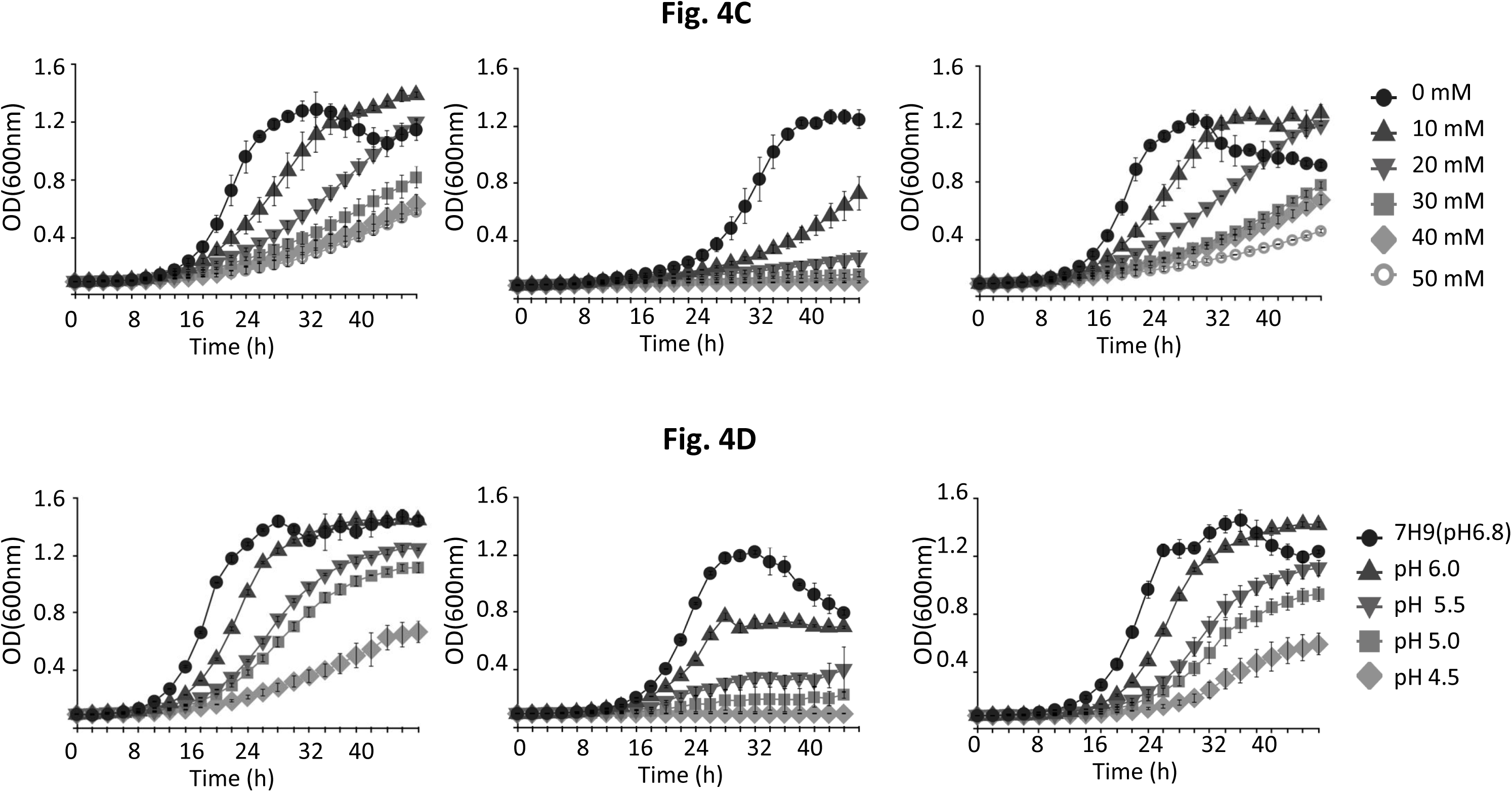
Growth of *Mycobacterium smegmatis* strains under stress conditions. Growth curve of Ms-WT (left panels), Ms-ΔRsmE (middle panels) and Ms-ΔRsmE/RsmE (right panels) with varying levels of (A) H_2_O_2_, (B) NaCl, (C) Sodium nitroprusside and (D) pH are plotted.

The growth of Ms-ΔRsmE as a function of decreasing pH, however, was more dramatic. Ms-WT showed slower growth as pH of the media was varied from pH 6.8 to pH 4.5. However growth of Ms-ΔRsmE, at pH 5.5 or lower was severely compromised compared to the Ms-WT cells (Figure 4D). The growth was restored similar to Ms-WT in the complemented Ms-ΔRsmE/RsmE strain, suggesting that expressing the gene in the knockout can reverse the sensitivity towards acidic stress (Figure 4D, right panel).

Taken together, these results highlight a possible role of RsmE in the acidic stress that mycobacteria might encounter during infection.

### Methylation at U1498 can modulate antibiotic response

U1498 is present in helix 44 of 30S subunit and is part of the functional unit, namely, P-site, of ribosome. Aminoglycosides and tetracyclines are the two major classes of drugs that bind at the functional site of 30S subunit of ribosome and were tested for drug susceptibility.

Role of m^3^ U1498 methylation in modulating drug response in mycobacteria was monitored by drug susceptibility assays of Ms-WT, Ms-ΔRsmE and Ms-ΔRsmE/RsmE (or Ms-ΔRsmE/RsmE’) (Figure 5). Growth in presence of kanamycin shows that a two fold higher concentration is required for growth inhibition of Ms-ΔRsmE as compared to Ms-WT (Figure 5A). In the MSMEG_4502 expressing complement strain, the growth inhibition was restored to similar kanamycin concentrations as Ms-WT, suggesting the difference of growth is associated with the methylation at U1498 (Figure 5A, right panel).

**Figure 5:**
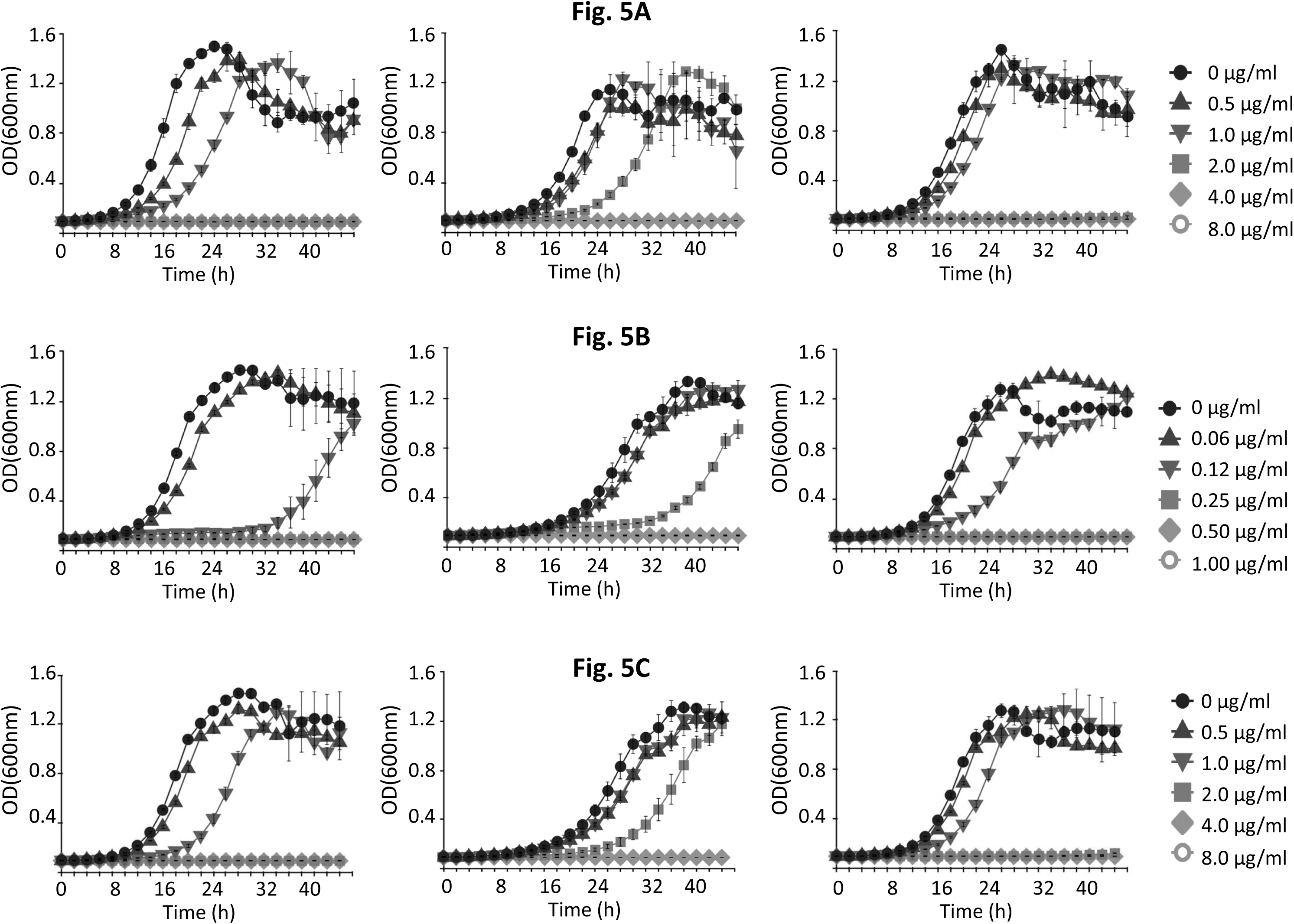
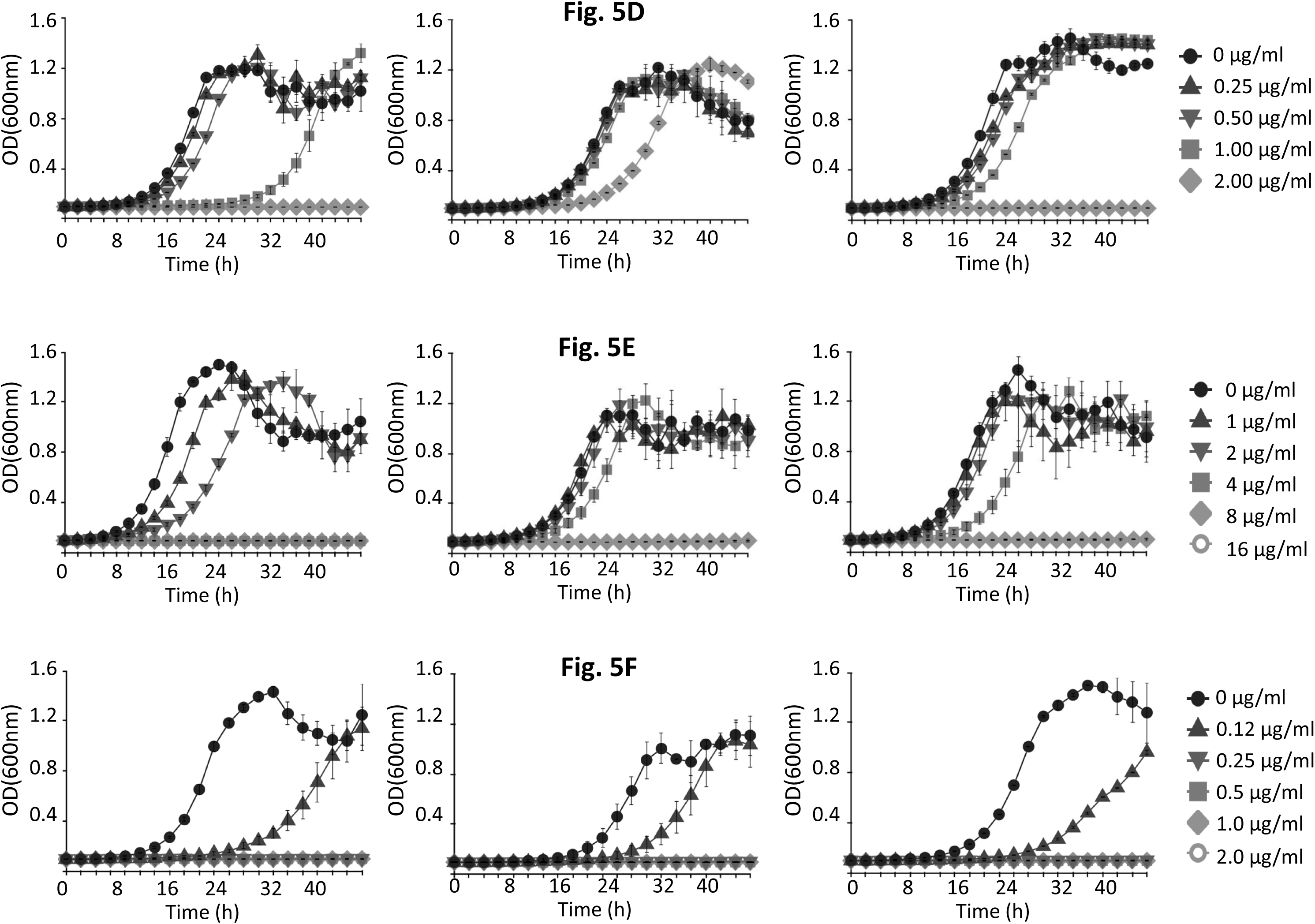
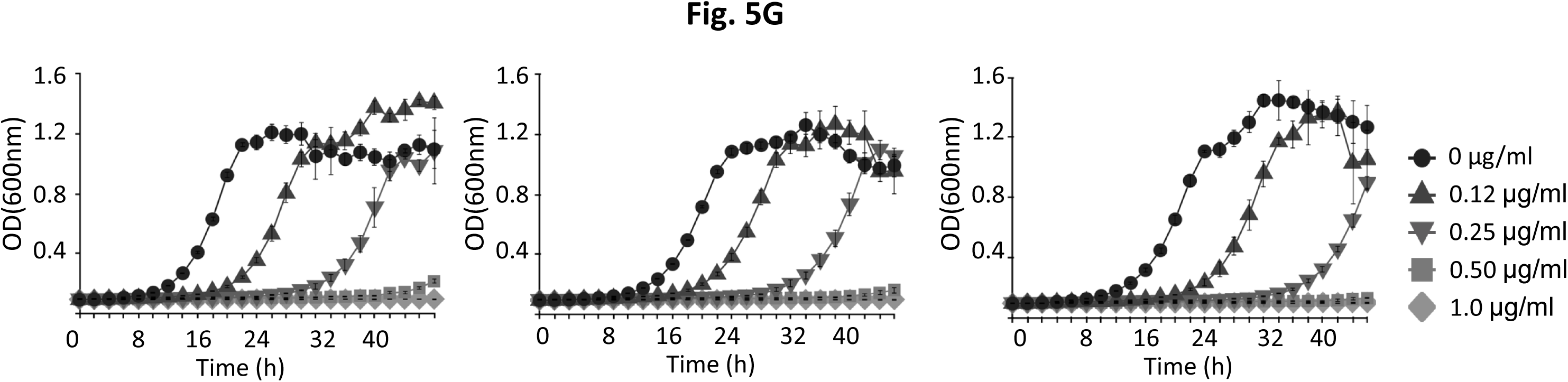
Drug response towards 30S targeting antibiotics. Growth of Ms-WT (left panels), Ms-ΔRsmE (middle panels) and complemented Ms-ΔRsmE/RsmE (or Ms-ΔRsmE/RsmE’), (right panels) were monitored with increasing concentrations of (A) kanamycin, (B) amikacin, (C) gentamicin, (D) paromomycin, (E) neomycin, (F) streptomycin and (G) tetracycline.

At least two fold higher concentration of drugs for growth inhibition is also required with amikacin (Figure 5B), gentamicin (Figure 5C) and paromomycin (Figure 5D) as compared to Ms-WT, which are restored in the Ms-ΔRsmE/RsmE complemented strain. Although a two fold higher concentration for growth inhibition is also required with neomycin (Figure 5E), it was not restored in the complemented strain.

The growth response of Ms-WT, Ms-ΔRsmE, Ms-ΔRsmE/RsmE with streptomycin (Figure 5F) and tetracycline (Figure 5G), which bind to A-site of the 30S subunit, farther from helix 44, however, showed no change in inhibitory concentrations.

Deletion of the RsmE-homolog of *M. smegmatis* hence leads to a low level of resistance to all the tested aminoglycosides that bind in the decoding center proximal to U1498 in helix 44 of 30S subunit of ribosome. The susceptibility is restored to lower concentrations of each aminoglycoside in the respective complemented (Ms-ΔRsmE/RsmE) strain in all cases (except neomycin), suggesting a correlation of methylation at U1498 with drug binding and antibiotic response for aminoglycosides that bind at this site.

### Antibiotic response to non-30S targeting drugs

To confirm whether the observed low level resistance was specific to aminoglycosides in Ms-ΔRsmE or did the deleted strain exhibit cross-resistance to any other antibiotics, growth of Ms-WT, Ms-ΔRsmE and Ms-ΔRsmE/RsmE was monitored with non-30S targeting drugs, namely; non-ribosomal drugs (rifampicin and moxiflacin) and 50S-targeting drugs (erythromycin and chloramphenicol). Growth of Ms-ΔRsmE with rifampicin (Figure 6A) that targets RNA-polymerase and with moxifloxacin (Figure 6B) that targets DNA gyrase was similar to that of Ms-WT and the inhibitory concentration for Ms-WT and Ms-ΔRsmE was same for both the drugs.

**Figure 6:**
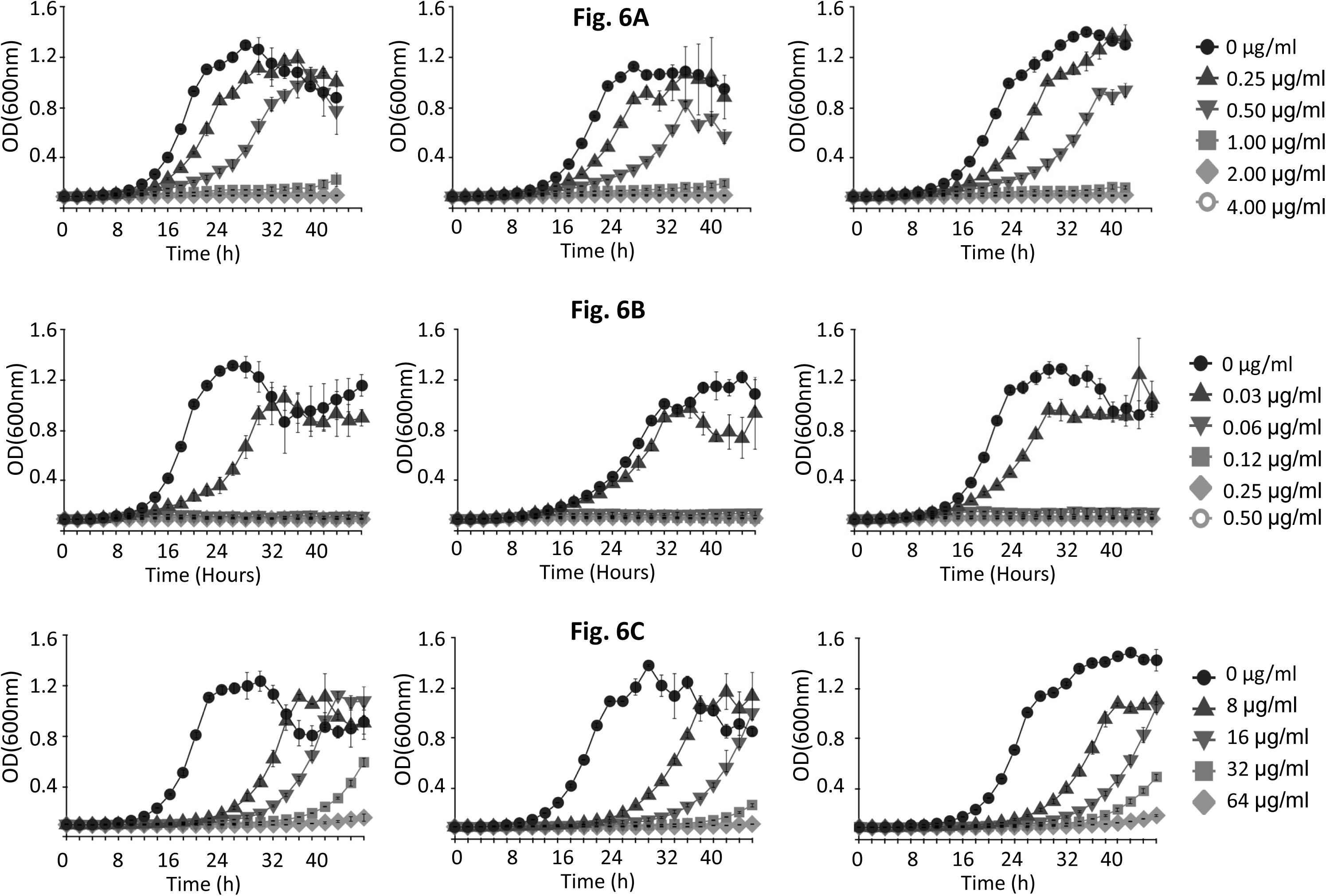
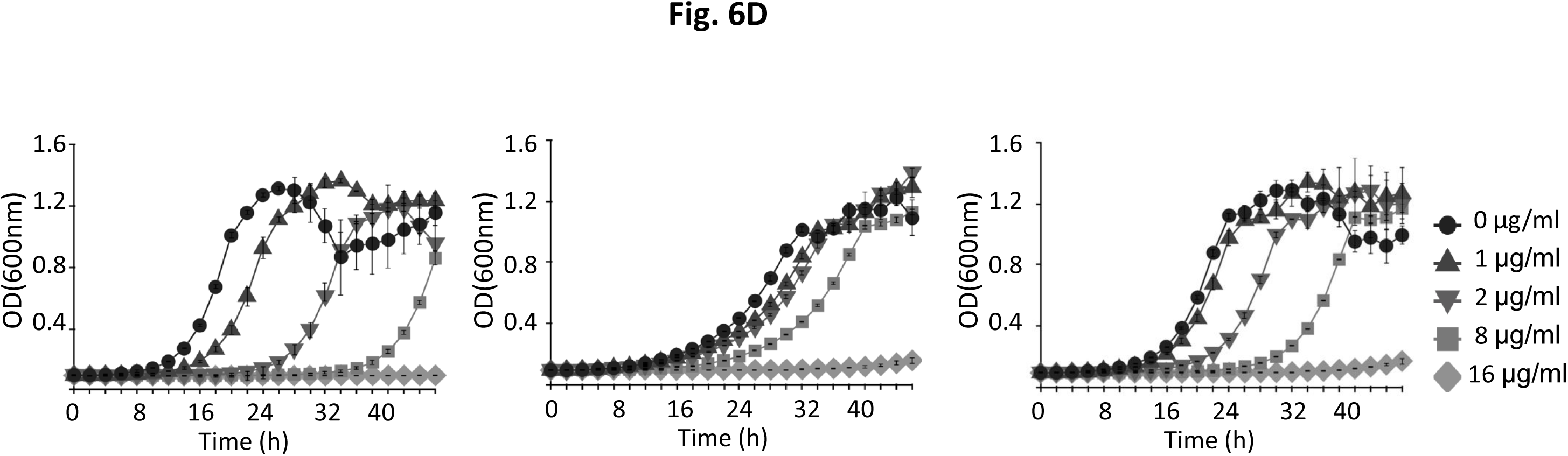
Drug response towards non-30S targeting antibiotics. Growth of Ms-WT (left panels), Ms-ΔRsmE (middle panels) and Ms-ΔRsmE/RsmE (right panels) were monitored with increasing concentrations of (A) rifampicin, (B) moxifloxacin, (C) erythromycin and (D) chloramphenicol.

Growth of Ms-ΔRsmE as compared to Ms-WT was found to be marginally affected by other drugs. Growth was slightly slower in response to erythromycin, that targets 50S subunit at the exit tunnel (Wilson 2014) (Figure 6C) but slightly faster in response to chloramphenicol, that targets ribosomal A-site in the 50S subunit (Wilson 2014) (Figure 6D). However, no change in inhibitory concentration was observed as compared to Ms-WT for both the drugs. Growth of Ms-ΔRsmE/RsmE complemented strain was similar to that of Ms-WT for all the non-30S targeting drugs (Figures 6, right panels).

Hence, the change in inhibitory concentrations, in the knockout strain of *M. smegmatis* (Ms-ΔRsmE) was specific towards antibiotics belonging primarily to aminoglycoside class among the tested drugs and does not show any cross-resistance to drugs of other classes (Table 2).

**Table 2:** Observed MICs for *M. smegmatis* strains

| Antibiotic | Class of antibiotic | Target of antibiotic | Ms-WT MIC | Ms-ΔRsmE MIC | Ms-ΔRsmE/RsmE MIC |
| --- | --- | --- | --- | --- | --- |
| **A. 30S targeting drugs** |  |  |  |  |  |
| Kanamycin | Aminoglycosides | 30S ribosome | 2.0 µg/ml | **4.0 µg/ml** | 2.0 µg/ml |
| Amikacin | Aminoglycosides | 30S ribosome | 0.25 µg/ml | **0.5 µg/ml** | 0.25 µg/ml |
| Gentamicin | Aminoglycosides | 30S ribosome | 2.0 µg/ml | **4.0 µg/ml** | 2.0 µg/ml |
| Neomycin | Aminoglycosides | 30S ribosome | 4.0 µg/ml | **8.0 µg/ml** | **8.0 µg/ml** |
| Paromomycin | Aminoglycosides | 30S ribosome | 2.0 µg/ml | **>4.0 µg/ml** | 2.0 µg/ml |
| Streptomycin | Aminoglycosides | 30S ribosome | 0.25 µg/ml | 0.25 µg/ml | 0.25 µg/ml |
| Tetracycline | Tetracyclines | 30S ribosome | 0.5 µg/ml | 0.5 µg/ml | 0.5 µg/ml |
| **B. 50S targeting drugs** |  |  |  |  |  |
| Erythromycin | Macrolides | 50S ribosome | 64.0 µg/ml | 64.0 µg/ml | 64.0 µg/ml |
| Chloramphenicol | Chloramphenicols | 50S ribosome | 16.0 µg/ml | 16.0 µg/ml | 16.0 µg/ml |
| **C. Non-ribosomal targeting drugs** |  |  |  |  |  |
| Moxifloxacin | Quinolones | DNA gyrase | 0.06 µg/ml | 0.06 µg/ml | 0.06 µg/ml |
| Rifampicin | Rifamycins | DNA<br>dependent<br>RNA<br>polymerase | 1.0 µg/ml | 1.0 µg/ml | 1.0 µg/ml |

### RsmE mutations are prevalent in the clinical strains of *M. tuberculosis*

Rv1010, Rv1407, Rv2966c, Rv2372c, Rv3919c, Rv2165 and Rv1003 were identified as homologs of conserved 16S rRNA methyltransferases RsmA, RsmB, RsmD, RsmE, RsmG, RsmH and RsmI, respectively, in 9745 clinical strains of *M. tuberculosis* in the PATRIC database (Wattam *et al.*, 2017). Non-synonymous mutations in the mycobacterial 16S rRNA methyltransferases were identified from genome data of all the strains, with *M. tuberculosis* H37Rv as the reference strain. Among all methyltransferases, RsmG, which is known to be associated with streptomycin resistance, has mutations in >40% of all strains. Surprisingly, at 19% of all strains with mutations, the RsmE homolog of *M. tuberculosis*, Rv2372c stands at second position, suggesting a key association of this methyltransferase with clinical manifestation and/or drug response in mycobacteria. Strains with mutations in RsmA, RsmH or RsmI comprised less than 3% of the strains, in each case, while mycobacterial strains carrying mutations in RsmD (Rv2966c) were negligible (Figure 7).

**Figure 7:**
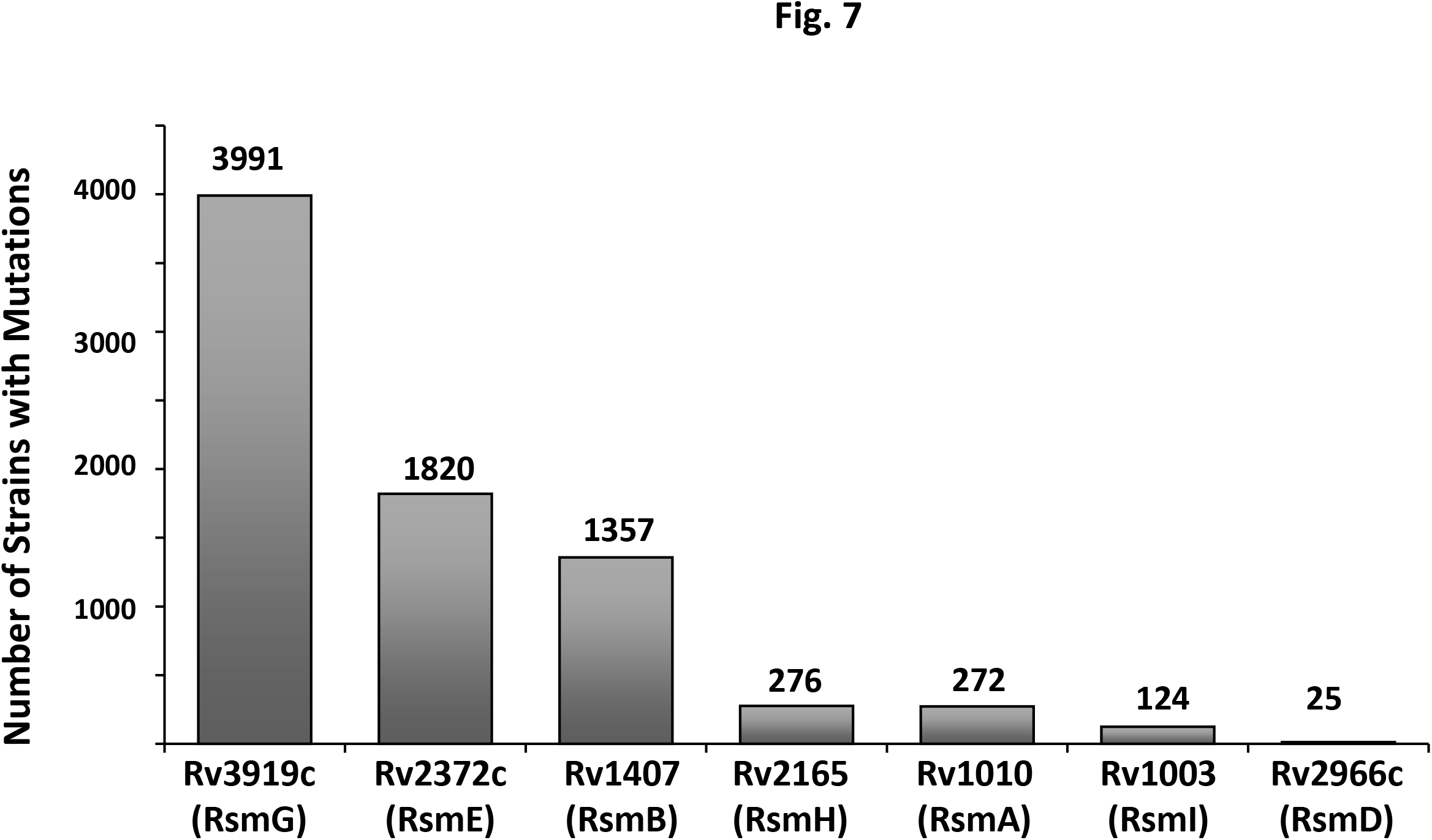
Mutations in 16S rRNA methyltransferases in clinical strains of *Mycobacterium tuberculosis*. The total number of non-synonymous mutations in the coding sequence of the conserved rRNA methyltransferases in the clinical strains of *M. tuberculosis* obtained from the PATRIC database is indicated. RsmG, having the highest number of mutations, is known to be associated with streptomycin resistance. RsmE methyltransferase has the second highest mutations among conserved methyltransferases, in a total of 19% clinical strains. The conserved *E. coli* counterpart for each mapped methyltransferase is also indicated.

## Discussion

U1498 is localized in the highly conserved region in the top half of helix 44 of 16S rRNA (Schuwirth *et al.*, 2005; Basturea 2006). This region is of high functional importance as it is part of the decoding centre. The decoding centre of ribosome ensures error-free selection of cognate tRNA to mRNA, promoting translation accuracy (Ogle *et al.*, 2003; Mahto and Chow 2013). Antibiotics of several classes, target the decoding centre by binding around helix 44 and inhibiting protein translation (Wilson 2014; Lin *et al.*, 2018).

Here, in this study, we have shown that methylation of U1498 of 16S rRNA is associated with altered antibiotic response of mycobacterial cells. This response is highly specific towards certain aminoglycosides targeting the 30S subunit of the ribosome. The binding site of these aminoglycosides in the ribosome was mapped from the available crystal structures in the PDB (Figure 8). Kanamycin, amikacin, neomicin, gentamicin and paromomycin belong to the class of antibiotics that bind near the top half of the helix 44, close to U1498 in 16S rRNA, whereas streptomycin and tetracycline bind at sites farther away. The fact that only the antibiotics that bind in the vicinity of U1498 exhibited altered growth response of Ms-ΔRsmE in presence of these aminoglycosides, suggests the absence of methylation in the deleted strain affects antibiotic binding at this site. A possible change in the local conformation of the binding site in the absence of m^3^U1498 methylation, might lead to change in the binding pattern of antibiotics, resulting in a low level increased resistance in Ms-ΔRsmE. This is the first report where methylation at U1498 of 16S rRNA is shown to be directly associated with change in the antibiotic response in mycobacteria.

**Figure 8:**
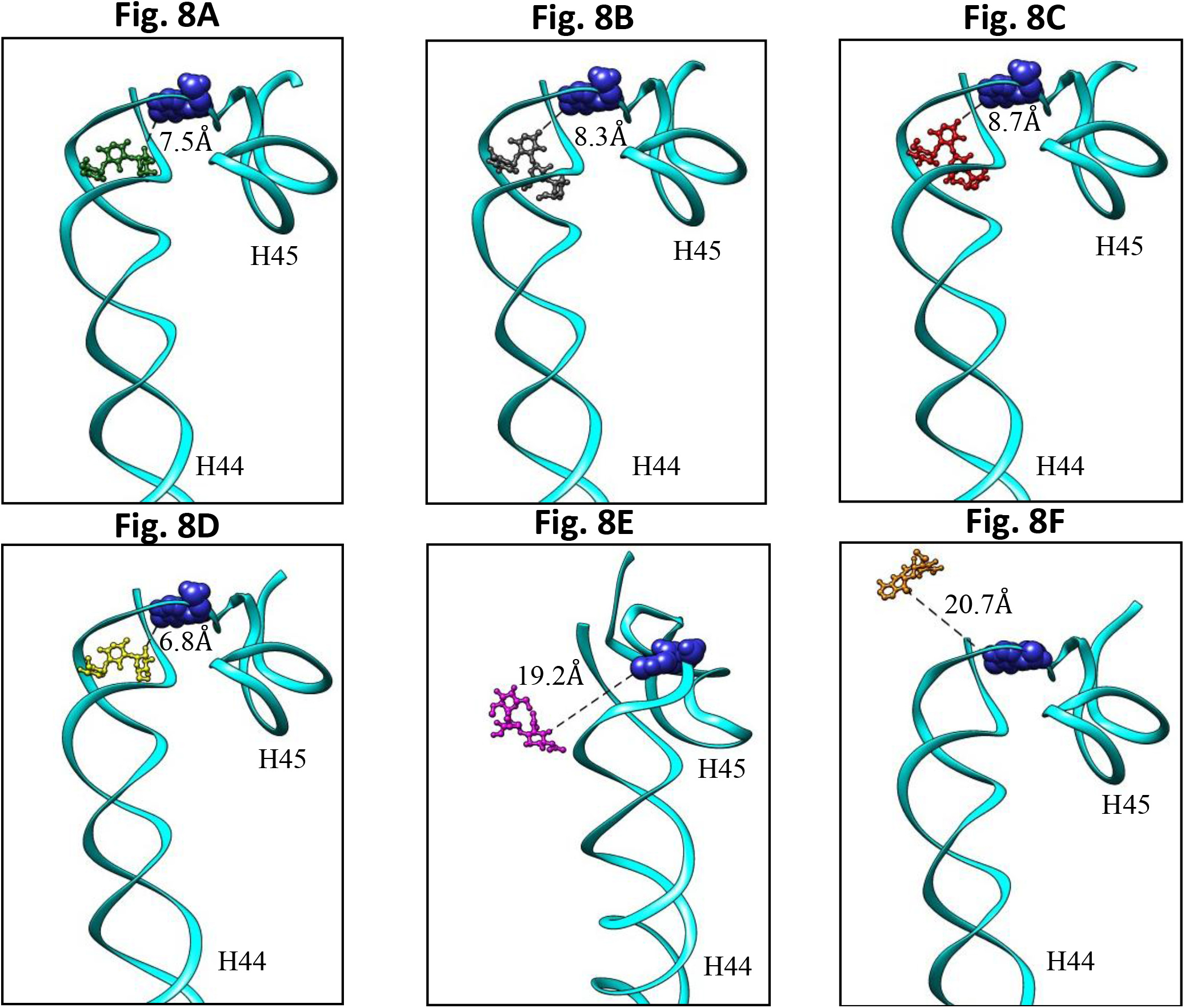
Binding site of aminoglycosides in ribosome strucutre. The binding of antibiotics (A) Gentamicin (green), (PDBID: 4V53), (B) paromomycin (grey), (PDBID: 4V5Y), (C) neomycin (red), (PDBID: 4V52), (D) tobramycin^a^ (yellow), (PDBID: 4LFC), (E) streptomycin (magenta), (PDBID: 4JI8) and (F) tetracycline (orange), (PDBID: 5J5B) was mapped using the available crystal structures in PDB and are shown as ball-and-stick structures. The distance from N-3 of U1498 to the bound drug (closest atom) is indicated in dashed lines (in Å). For clarity, only helix 44 and helix 45 of 30S ribosomal subunit are shown here (cyan). U1498 is indicated as space-filled structure (bright blue). Non-30S targeting drugs are not shown as they bind much farther away from U1498. a: Tobramycin-bound structure is indicative of kanamycin/amikacin binding site, as a kanamycin/amikacin-bound ribosome structure could not be identified in PDB.

RsmE is widely distributed in major bacterial groups (Baldridge and Contreras 2014; Benítez-Páez *et al.*, 2014). Despite a reductive evolution of the mycobacterial genomes, conservation of Rv2372c, the RsmE-homolog in *M. tuberculosis*, suggests m^3^U1498 methylation is necessary for normal cellular physiology. The m^3^U1498 nucleotide interacts directly with the mRNA in the ternary complex of 70S ribosome-mRNA-tRNA of *Thermus thermophilus* (Korostelev *et al.*, 2006) and *E. coli* (Selmer 2006) highlighting its role in efficient translation. Moreover, a U1498G mutation blocks the first peptide bond formation (Cunningham *et al.*, 1993) due to possible change in tertiary structure around the decoding structure (Ringquist *et al.*, 1993). The crystal structure of 70S ribosome also showed that m^3^U1498 is present in the intersubunit bridge of 50S and 30S and interacts with the conserved regions in 23S rRNA (Selmer 2006; Baldridge and Contreras 2014). Despite taking part in several key interactions in the assembly and function of ribosome, the RsmE knockout of *E. coli* grows normally although it fails to compete when grown together with the wild type (Basturea 2006). In case of *M. smegmatis*, although the RsmE-homolog, *MSMEG_4502*, is not essential for its survival, knockout of *MSMEG_4502* leads to subtle changes in physiology of the mutant Ms-ΔRsmE strain.

Even a low level resistance in a microorganism *in vitro*, can serve as a stepping stone to development of high level clinically relevant resistant strains (Gomez *et al.*, 2017), as seen for streptomycin (Okamoto *et al.*, 2007). The modulation of antibiotic response by RsmE methyltransferase might lead to a potential threat and hence must be understood in greater details. Mycobacteria encounter a wide variety of stress, *viz.*, oxidative and pH stress during infection of the host. In a recent report, phagosomal acidification was shown to alter the redox physiology of *M. tuberculosis* to promote tolerance of the bacterium to antibiotics (Mishra *et al.*, 2019). Inhibition of phagosomal acidification by the antimalarial drug chloroquine eradicated drug-tolerant bacilli in *in vivo* models. The increased sensitivity of Ms-ΔRsmE, lacking methylation at 1498 position, to acid stress, however, suggests an alternate potential combinatorial way to target tubercular bacilli and prevent emergence of resistance to front line drugs.

Overall, we show that absence of methylation of U1498 of 16S rRNA leads to development of low level resistance in *M. smegmatis*. The increased resistance was specific only for some antibiotics belonging to the aminoglycoside class. Mapping of binding site in the ribosome, showed that growth of Ms-ΔRsmE was affected specifically by aminoglycosides that bind in the vicinity of U1498 of 16S rRNA. We also found that RsmE has the second highest mutation rate in the clinical strains among all conserved methyltransferases. This reinforces the possibility of association of RsmE towards development of resistance in clinical strains of*M. tuberculosis*, which must be tackled in a timely manner.

## Acknowledgements

SB acknowledges Senior Research Fellowship from CSIR. The authors thank Anjali Maurya and Pooja Badhwar for useful discussions. We thank CSIR-Institute of Genomics and Integrative Biology for providing the infrastructure and central instrument lab facility. BT acknowledges financial support from Department of Science and Technology (DST) (EMR/2016/003589) Government of India. pTC-MCS and pTC0X1L were kind gifts from Prof. Sabine Ehrt, Weill Cornell Medical College; pMSG360zeo was a kind gift from Prof. Michael Glickman, Memorial Sloan Kettering Cancer Center and is gratefully acknowledged.

## References

Baldridge, Kevin C. and Lydia M. Contreras. 2014. “Functional Implications of Ribosomal RNA Methylation in Response to Environmental Stress.” Critical Reviews in Biochemistry and Molecular Biology 49(1):69–89.

Barkan, Daniel, Vivek Rao, George D. Sukenick, and Michael S. Glickman. 2010. “Redundant Function of CmaA2 and MmaA2 in *M. tuberculosis* Cis Cyclopropanation of Oxygenated Mycolates.” Journal of Bacteriology 192(14):3661–68.

Basturea, G. N. 2006. “Identification and Characterization of RsmE, the Founding Member of a New RNA Base Methyltransferase Family.” RNA 12(3):426–34.

Benítez-Páez, Alfonso, Sonia Cárdenas-Brito, Mauricio Corredor, Magda Villarroya, and María Eugenia Armengod. 2014. “Impairing Methylationcs at Ribosome RNA, a Point Mutation-Dependent Strategy for Aminoglycoside Resistance: The RsmG Case.” Biomedica 34(SUPPL.1):41–49.

Borovinskaya, Maria A., Raj D. Pai, Wen Zhang, Barbara S. Schuwirth, James M. Holton, Go Hirokawa, Hideko Kaji, Akira Kaji, and Jamie H. Doudna Cate. 2007. “Structural Basis for Aminoglycoside Inhibition of Bacterial Ribosome Recycling.” Nature Structural & Molecular Biology 14(8):727–32.

Borovinskaya, Maria A., Shinichiro Shoji, Kurt Fredrick, and Jamie H. D. Cate. 2008. “Structural Basis for Hygromycin B Inhibition of Protein Biosynthesis.” RNA 14(8):1590–99.

Buriánková, Karolína, Florence Doucet-Populaire, Olivier Dorson, Anne Gondran, Jean Claude Ghnassia, Jaroslav Weiser, and Jean Luc Pernodet. 2004. “Molecular Basis of Intrinsic Macrolide Resistance in the *M. tuberculosis* Complex.” Antimicrobial Agents and Chemotherapy.

Cocozaki, Alexis I., Roger B. Altman, Jian Huang, Ed T. Buurman, Steven L. Kazmirski, Peter Doig, D. Bryan Prince, Scott C. Blanchard, Jamie H. D. Cate, and Andrew D. Ferguson. 2016. “Resistance Mutations Generate Divergent Antibiotic Susceptibility Profiles against Translation Inhibitors.” Proceedings of the National Academy of Sciences of the United States of America.

Cunningham, Philip R., Kelvin Nurse, Carl J. Weitzmann, and James Ofengand. 1993. “Functional Effects of Base Changes Which Further Define the Decoding Center of Escherichia Coli 16S Ribosomal RNA: Mutation of C1404, G1405, C1496, G1497, and U1498.” Biochemistry 32(28):7172–80.

Demirci, Hasan, Leyi Wang, F. V. Murphy, Eileen L. Murphy, Jennifer F. Carr, Scott C. Blanchard, Gerwald Jogl, Albert E. Dahlberg, and Steven T. Gregory. 2013. “The Central Role of Protein S12 in Organizing the Structure of the Decoding Site of the Ribosome.” RNA 19(12):1791–1801.

Finken, Marion, Philip Kirschner, Albrecht Meier, Annette Wrede, and Erik C. Böttger. 1993. “Molecular Basis of Streptomycin Resistance in *M. tuberculosis:* Alterations of the Ribosomal Protein S12 Gene and Point Mutations within a Functional 16S Ribosomal RNA Pseudoknot.” Molecular Microbiology 9(6):1239–46.

Gaora, Peadar, Simona Barnini, Chris Hayward, Elaine Filley, Graham Rook, Douglas Young, and Jelle Thole. 1997. “Mycobacteria as Immunogens: Development of Expression Vectors for Use in Multiple Mycobacterial Species.” Medical Principles and Practice.

Golovina, Anna Y., Margarita M. Dzama, Ilya a Osterman, Petr V Sergiev, Marina V Serebryakova, Alexey a Bogdanov, and Olga a Dontsova. 2012. “The Last RRNA Methyltransferase of E. Coli Revealed: The YhiR Gene Encodes Adenine-N6 Methyltransferase Specific for Modification of A2030 of 23S Ribosomal RNA.” RNA (New York, N.Y.) 18(9):1725–34.

Gomez, James E., Benjamin B. Kaufmann-Malaga, Carl N. Wivagg, Peter B. Kim, Melanie R. Silvis, Nikolai Renedo, Thomas R. Ioerger, Rushdy Ahmad, Jonathan Livny, Skye Fishbein, James C. Sacchettini, Steven A. Carr, and Deborah T. Hung. 2017. “Ribosomal Mutations Promote the Evolution of Antibiotic Resistance in a Multidrug Environment.” ELife 6.

Guo, Xinzheng V, Mercedes Monteleone, Marcus Klotzsche, Annette Kamionka, Wolfgang Hillen, Miriam Braunstein, Sabine Ehrt, and Dirk Schnappinger. 2007. “Silencing Essential Protein Secretion in *M. smegmatis* by Using Tetracycline Repressors.” Journal of Bacteriology 189(13):4614–23.

Gygli, Sebastian M., Sonia Borrell, Andrej Trauner, and Sebastien Gagneux. 2017. “Antimicrobial Resistance in *M. tuberculosis:* Mechanistic and Evolutionary Perspectives.” FEMS Microbiology Reviews.

Hameed, H. M. Adna., Md Mahmudul Islam, Chiranjibi Chhotaray, Changwei Wang, Yang Liu, Yaoju Tan, Xinjie Li, Shouyong Tan, Vincent Delorme, Wing W. Yew, Jianxiong Liu, and Tianyu Zhang. 2018. “Molecular Targets Related Drug Resistance Mechanisms in MDR-, XDR-, and TDR-*M. tuberculosis* Strains.” Frontiers in Cellular and Infection Microbiology 8(APR).

Kambli, Priti, Kanchan Ajbani, Chaitali Nikam, Meeta Sadani, Anjali Shetty, Sophia B. Georghiou, Timothy C. Rodwell, Antonino Catanzaro, P. D. Hinduja National Hospital, P. D. Hinduja National Hospital, San Diego, and San Diego. 2016. “Correlating Rrs and Eis Promoter Mutations in Clinical Isolates of M. tuberculosis with Phenotypic Susceptibility Levels to the Second Line Injectables.” 5(1):1–6.

Klotzsche, Marcus, Sabine Ehrt, and Dirk Schnappinger. 2009. “Improved Tetracycline Repressors for Gene Silencing in Mycobacteria.” Nucleic Acids Research 37(6):1778–88.

Korostelev, Andrei, Sergei Trakhanov, Martin Laurberg, and Harry F. Noller. 2006. “Crystal Structure of a 70S Ribosome-TRNA Complex Reveals Functional Interactions and Rearrangements.” Cell 126(6):1065–77.

Kumar, Atul, Santosh Kumar, and Bhupesh Taneja. 2014. “The Structure of Rv2372c Identifies an RsmE-like Methyltransferase from *M. tuberculosis*” Acta Crystallographica Section D: Biological Crystallography 70(3):821–32.

Kumar, Atul, Kashyap Saigal, Ketan Malhotra, Krishna Murari Sinha, and Bhupesh Taneja. 2011. “Structural and Functional Characterization of Rv2966c Protein Reveals an RsmD-like Methyltransferase from *M. tuberculosis* and the Role of Its N-Terminal Domain in Target Recognition.” Journal of Biological Chemistry 286(22):19652–61.

Lin, Jinzhong, Dejian Zhou, Thomas A. Steitz, Yury S. Polikanov, and Matthieu G. Gagnon. 2018. “Ribosome-Targeting Antibiotics: Modes of Action, Mechanisms of Resistance, and Implications for Drug Design.” Annual Review of Biochemistry 87(1):451–78.

Louw, G. E., R. M. Warren, N. C. Gey Van Pittius, C. R. E. McEvoy, P. D. Van Helden, and T. C. Victor. 2009. “A Balancing Act: Efflux/Influx in Mycobacterial Drug Resistance.” Antimicrobial Agents and Chemotherapy 53(8):3181–89.

Mahto, Santosh K. and Christine S. Chow. 2013. “Probing the Stabilizing Effects of Modified Nucleotides in the Bacterial Decoding Region of 16S Ribosomal RNA.” Bioorganic and Medicinal Chemistry 21(10):2720–26.

Maus, Courtney E., Bonnie B. Plikaytis, and Thomas M. Shinnick. 2005. “Mutation of TlyA Confers Capreomycin Resistance in *M. tuberculosis*.” Antimicrobial Agents and Chemotherapy.

Mishra, Richa, Sakshi Kohli, Nitish Malhotra, Parijat Bandyopadhyay, Mansi Mehta, MohamedHusen Munshi, Vasista Adiga, Vijay Kamal Ahuja, Radha K. Shandil, Raju S. Rajmani, Aswin Sai Narain Seshasayee, and Amit Singh. 2019. “Targeting Redox Heterogeneity to Counteract Drug Tolerance in Replicating *M. tuberculosis*.” Science Translational Medicine 11(518):eaaw6635.

Nosova, Elena Yu, Anastasia A. Bukatina, Yulia D. Isaeva, Marina V. Makarova, K. Y. Galkina, and Arkadyi M. Moroz. 2013. “Analysis of Mutations in the GyrA and GyrB Genes and Their Association with the Resistance of *M. tuberculosis* to Levofloxacin, Moxifloxacin and Gatifloxacin.” Journal of Medical Microbiology 62(Pt_1):108–13.

Ogle, James M., Andrew P. Carter, and V. Ramakrishnan. 2003. “Insights into the Decoding Mechanism from Recent Ribosome Structures.” Trends in Biochemical Sciences 28(5):259–66.

Okamoto, Susumu, Aki Tamaru, Chie Nakajima, Kenji Nishimura, Yukinori Tanaka, Shinji Tokuyama, Yasuhiko Suzuki, and Kozo Ochi. 2007. “Loss of a Conserved 7-Methylguanosine Modification in 16S RRNA Confers Low-Level Streptomycin Resistance in Bacteria.” Molecular Microbiology 63(4):1096–1106.

Pelicic, Vladimir, Mary Jackson, J. M. Reyrat, William R. Jacobs, Brigitte Gicquel, and Christophe Guilhot. 1997. “Efficient Allelic Exchange and Transposon Mutagenesis in M. tuberculosis Proceedings of the National Academy of Sciences 94(20):10955–60.

Pelicic, Vladimir, Jean-Marc Reyrat, and Brigitte Gicquel. 1996. “Generation of Unmarked Directed Mutations in Mycobacteria, Using Sucrose Counter-Selectable Suicide Vectors.” Molecular Microbiology 20(5):919–25.

Pfister, P., M. Risch, D. E. Brodersen, and E. C. Bottger. 2003. “Role of 16S RRNA Helix 44 in Ribosomal Resistance to Hygromycin B.” Antimicrobial Agents and Chemotherapy 47(5):1496–1502.

Reeves, Analise Z., Patricia J. Campbell, Razvan Sultana, Seidu Malik, Megan Murray, Bonnie B. Plikaytis, Thomas M. Shinnick, and James E. Posey. 2013. “Aminoglycoside Cross-Resistance in *M. tuberculosis* Due to Mutations in the 5’ Untranslated Region of WhiB7.” Antimicrobial Agents and Chemotherapy 57(4):1857–65.

Ringquist, Steven, Philip Cunningham, Carl Weitzmann, Leo Formenoy, Cornelius Pleij, James Ofengand, and Larry Gold. 1993. “Translation Initiation Complex Formation with 30 S Ribosomal Particles Mutated at Conserved Positions in the 3’-Minor Domain of 16 S RNA.” Journal of Molecular Biology 234(1):14–27.

Schuwirth, Barbara S., Maria A. Borovinskaya, Cathy W. Hau, Wen Zhang, Antón Vila-Sanjurjo, James M. Holton, and Jamie H. Doudna Cate. 2005. “Structures of the Bacterial Ribosome at 3.5 Å Resolution.” Science 310(5749):827–34.

Selmer, Maria. 2006. “Structure of the 70S Ribosome Complexed with MRNA and TRNA.” Science 313(5795):1935–42.

Sotgiu, Giovanni, Rosella Centis, L. D’ambrosio, and G. B. Migliori. 2015. “Tuberculosis Treatment and Drug Regimens.” Cold Spring Harbor Perspectives in Medicine 5(5):a017822– a017822.

Sowajassatakul, Angkanang, Therdsak Prammananan, Angkana Chaiprasert, and Saranya Phunpruch. 2014. “Molecular Characterization of Amikacin, Kanamycin and Capreomycin Resistance in M/XDR-TB Strains Isolated in Thailand.” BMC Microbiology 14(1):1–7.

Wattam, Alice R., James J. Davis, Rida Assaf, Sébastien Boisvert, Thomas Brettin, Christopher Bun, Neal Conrad, Emily M. Dietrich, Terry Disz, Joseph L. Gabbard, Svetlana Gerdes, Christopher S. Henry, Ronald W. Kenyon, Dustin Machi, Chunhong Mao, Eric K. Nordberg, Gary J. Olsen, Daniel E. Murphy-Olson, Robert Olson, Ross Overbeek, Bruce Parrello, Gordon D. Pusch, Maulik Shukla, Veronika Vonstein, Andrew Warren, Fangfang Xia, Hyunseung Yoo, and Rick L. Stevens. 2017. “Improvements to PATRIC, the All-Bacterial Bioinformatics Database and Analysis Resource Center.” Nucleic Acids Research 45(D1):D535–42.

WHO. 2018. Global TB Report 2018.

Wilson, Daniel N. 2014. “Ribosome-Targeting Antibiotics and Mechanisms of Bacterial Resistance.” Nature Reviews Microbiology 12(1):35–48.

Wong, Sharon Y., Jong Seok Lee, Hyun Kyung Kwak, Laura E. Via, Helena I. M. Boshoff, and Clifton E. Barry. 2011. “Mutations in GidB Confer Low-Level Streptomycin Resistance in *M. tuberculosis*” Antimicrobial Agents and Chemotherapy 55(6):2515–22.

Zhang, Y. and W. W. Yew. 2015. “Mechanisms of Drug Resistance in *M. tuberculosis:* Update 2015.” International Journal of Tuberculosis and Lung Disease 19(11):1276–89.

